# Population-level constraints on single neuron tuning changes during behavioral adaptation

**DOI:** 10.1101/2025.04.17.649401

**Authors:** Hannah M. Stealey, Yi Zhao, Hung-Yun Lu, Enrique Contreras-Hernandez, Yin-Jui Chang, Philippe Tobler, Jarrod A. Lewis-Peacock, Samantha R. Santacruz

## Abstract

In neurofeedback training, select neurons – of the billions that comprise the brain – are able to adapt their activity to produce a desired behavior, even though the feedback signal is low-dimensional. However, the extent that individual neurons are able to alter their activity within the context of the neurofeedback population is unclear. Using a brain-computer interface in monkeys (n=2; *Macaca mulatta*), we tested how the tuning of individual neurons changed in response to various feedback perturbations. Overall, we found that neurons that co-varied their activity the most with the population and those that contributed most to the behavior prior to the perturbation were most resistant to change– regardless of perturbation type. The degree to which individual neurons counteracted the applied perturbation explained daily differences in how well neurons collectively produced adapted behavior. Our work provides important insight into how population-level constraints impact the ability of individual neurons to adapt and support behavioral flexibility.

## INTRODUCTION

Throughout life, the healthy brain maintains the remarkable ability to maintain robust connections, to form new connections, and to adapt connections when necessary to navigate successfully through an ever-changing environment^1–4^. This ability, termed “neuroplasticity”, can be purposefully harnessed through neurofeedback training. In neurofeedback training, individuals produce specific neural activity patterns to ultimately facilitate desired behavioral changes. Neurofeedback training involves translating (“decoding”) neural activity into an external perceptible (e.g., visual, auditory) signal through a mathematical mapping (“decoder”). Given the external representation of their internal neural signals, subjects can volitionally modulate their brain activity to produce a specific pattern of brain activity. While the clinical use of neurofeedback training has shown promise, it also has had vast clinical variability, limiting its full potential for widespread adoption^5,6^. Clinical variability is partly driven by non-standardized training protocols, some of which may be more difficult to learn^6^. Therefore, a deeper understanding of the limitations in neuronal adaptability and their impact on desired behavioral changes would improve clinical neurofeedback paradigms^7^.

Evidence as early as 1969^8^ has demonstrated that specific individual neurons *in vivo* can alter their activity in response to low-dimensional neurofeedback signals, despite being processed by the brain’s multi-billion-dimensional network. This finding has since inspired brain-computer interface (BCI) studies that have studied the relationship between neural activity changes and neurofeedback signals through investigating how subjects learn a novel decoder or adapt to a perturbation of a previously learned decoder^9–21^. These studies have demonstrated that neurons directly involved in behavioral output (i.e., potent neurons) change their activity more than non-potent neurons. Similarly, neurons that are impacted by changes to the decoder (i.e., perturbed neurons) also change their activity more non-perturbed neurons. Together, these results suggest that feedback causally influences neural activity but highlight that non-potent and non-perturbed neurons still alter their activity. However, the mechanisms through which this underlying network constrains changes in neural activity are not well understood, especially on the short timescale of behavioral adaptation^22,23^.

To address this gap, we analyzed individual neuron changes in the context of their relationship to the selected neurofeedback population (“BCI neurons”) during a BCI task. Specifically, we investigated how properties of the neural repertoire prior to the introduction of a feedback perturbation influenced the ability of individual neurons to adapt. Through perturbing neurofeedback signals in a BCI task, we assessed how a neuron’s task-relevancy and relationship to the BCI population impacted its ability to appropriately alter its activity. We then determined how well a neuron’s ability to adapt to varying difficulties of protocols translated to behavioral changes and explained behavioral differences.

## RESULTS

We previously^24^ trained monkeys (n=2; *Macaca mulatta*) in a common ^9,10,12,13,15,16,18,25^ 8-target, center-out BCI task in which they modulated spiking activity from cortical motor neurons to drive a computer cursor (**Figure 1A**). To mathematically map neural activity to cursor velocity, we implemented a Kalman filter decoder (**Figure 1A**, *see Methods – Kalman filter decoder*). In each center-out trial, subjects were required to move the cursor to a peripheral target within 10 seconds (**Figure 1B**). Following a block of baseline trials to establish a within-subject and within-session control, at the onset of the perturbation block of trials, we introduced one type of decoder perturbation: an easy rotation perturbation (50°), a hard rotation perturbation (90°), or a shuffle perturbation (**Figure 1B**). In general, the decoder perturbation impacted the Kalman gain (**K**) assigned to each neuron (for all neurons equally (rotation perturbation) or randomly for a subset of neurons (shuffle perturbation)). This alteration of gain remained fixed throughout the perturbation block of trials. To regain proficient control of the cursor (i.e., move the cursor towards the target), subjects needed to intentionally alter their neural activity to account for the difference between their intended movement under the baseline decoder and the perturbed feedback. Under the rotation perturbation, the cursor initially appeared rotated relative to the intended direction. Under the shuffle perturbation, the disruption to cursor movement was less predictable and varied by session.

**Figure 1.**
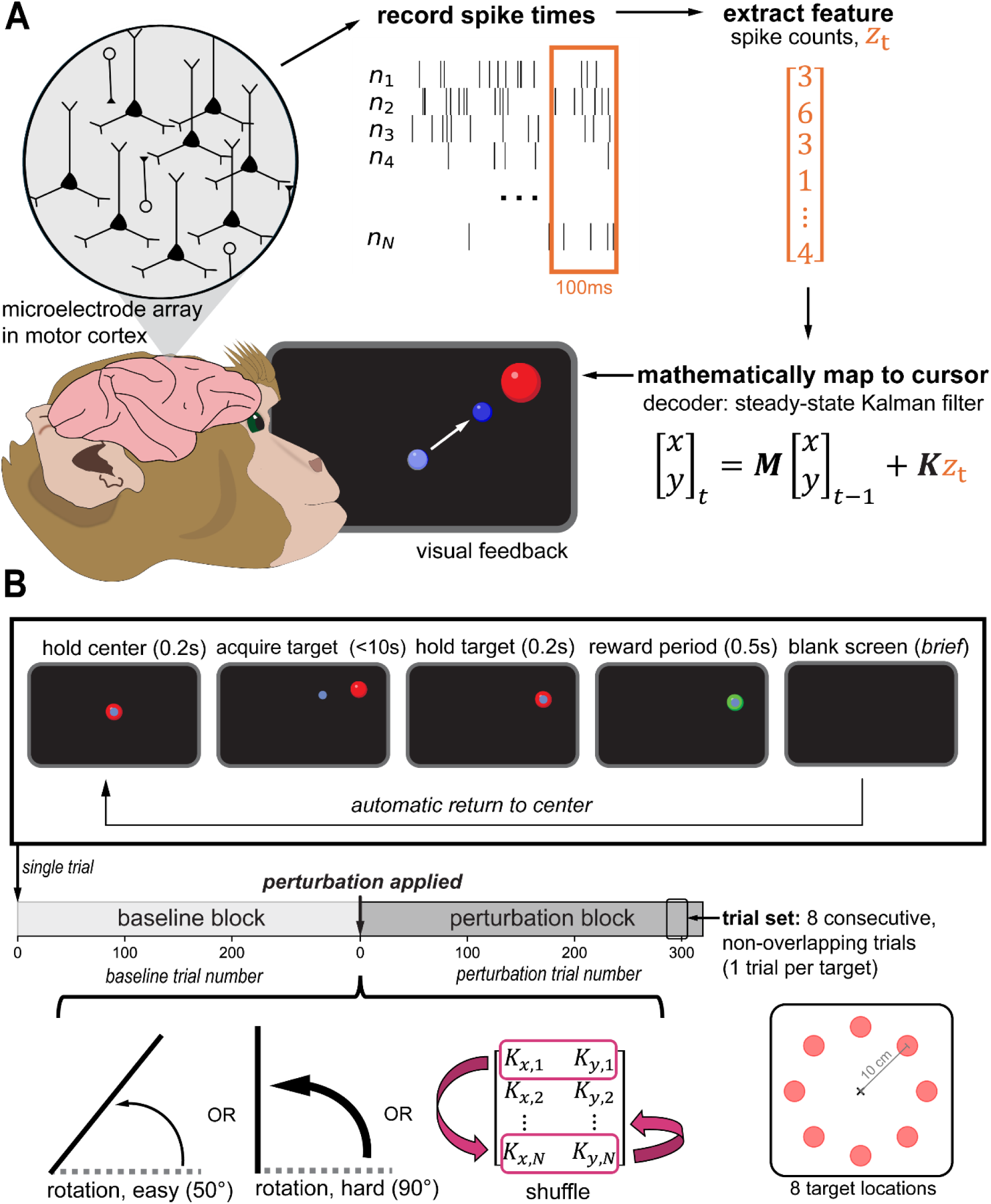
| Experimental design. (**A**) *BCI schematic.* Subjects modulated motor cortical (forelimb area of primary motor cortex (M1)/ pre-motor cortex (PMd)) spiking activity to control a computer cursor (blue, radius 0.5cm) to 1 of 8 possible peripheral targets (red). Neural activity was mapped to cursor velocity through a Kalman filter decoder. In the filter, spike count vectors (z_t_) collected every 100ms (10Hz) were weighted by a Kalman gain matrix (**K**) for x– and y– velocity components. This product was then added to a matrix of constants (**M**) projected a portion of the previous velocity state (*t-1*) to the current timestep (*t*). (**B**) *Single trial and single session structure.* Each trial followed the same series of steps. Subjects moved the cursor to the presented target within 10s. To ensure that the target acquisition was intentional, subjects were required to kept the cursor within the target bounds for 200ms to successfully complete a trial. Subjects completed a baseline block of trials and perturbation block of trials. Each block was divided into “trial sets” which contained one trial to each of the 8 target locations. At the onset of the perturbation block, 1 of 3 possible perturbation types were implemented on **K**: an easy rotation perturbation (50°), a hard rotation perturbation (90°), or a shuffle perturbation. The perturbation remained fixed for the duration of the perturbation block.

### Behavior under a shuffle perturbation does not exhibit typical adaptation characteristics

We first examined how subject behavior evolved over the course of the perturbation block in response to the different types of perturbation. Under both the rotation and shuffle perturbation types, cursor trajectories were typically direct-to-target and across baseline trials under both perturbation types (**Figure 2** –*first and second column*). Under the rotation condition, the trajectories in early perturbation trials were curved in proportion to the applied rotation and remain curved (but less so) during late perturbation trials (**Figure 2** –*third and fourth column, first row*). In contrast, the shuffle perturbation caused the cursor to move in a more unpredictable manner during early perturbation trials (**Figure 2** –*third column, second row*). Still, subjects were able to improve the directness of the cursor trajectories as demonstrated by the example late perturbation trial trajectories (**Figure 2** –*fourth column, second row*).

**Figure 2.**
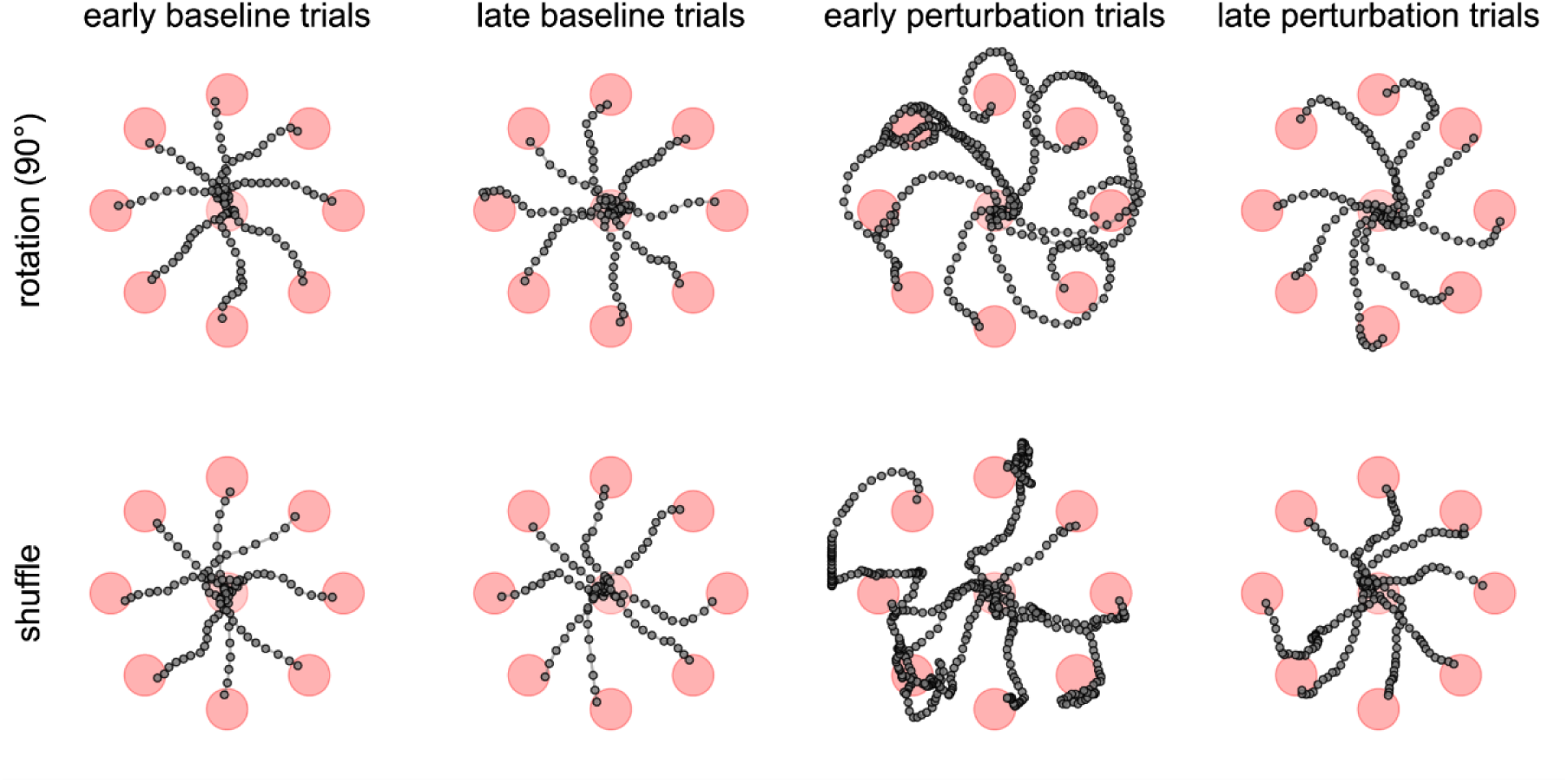
| Sample cursor trajectories under the rotation and shuffle perturbation (Monkey A). Each subplot contains superimposed views of cursor trajectories from one sample trial to each of the possible target locations (8) under the hard rotation perturbation (top row) and the shuffle perturbation (bottom row) during different stages of the task. During the task, subjects could only see the current position of the cursor.

We quantified these observations by assessing how trial time and cursor path length (“distance”) changed relative to average baseline performance over sets of perturbation trials (*Methods – Behavioral metrics*). Over the course of the perturbation block, behavioral performance followed an exponential decay (**Figure 3A**). Under both rotation conditions, performance initially increased rapidly (as indicated by decreases in trial times and distances relative to baseline) and quickly reached an asymptote with trial times still increased relative to baseline. Similarly, under the shuffle perturbation, performance was immediately negatively impacted following the onset of the shuffling (**Figure 3A**). What separates behavior under the two perturbation types is the subsequent temporal profiles. While performance improved exponentially under the rotation perturbation (i.e., values approached baseline), performance under the shuffle perturbation remained impacted, indicated by flat performance profiles (Monkey A: slope=-0.004%/trial set, p=0.90; Monkey B, slope=-0.07%/trial set, p=0.03; linear regression). While the performance under the rotation condition is consistent with classically-observed adaptation profiles (characteristics: rapid improvements and suboptimal adaptation in NHPs^10,11,26^, humans^27–37^), adaptation under the shuffle perturbation was not.

**Figure 3.**
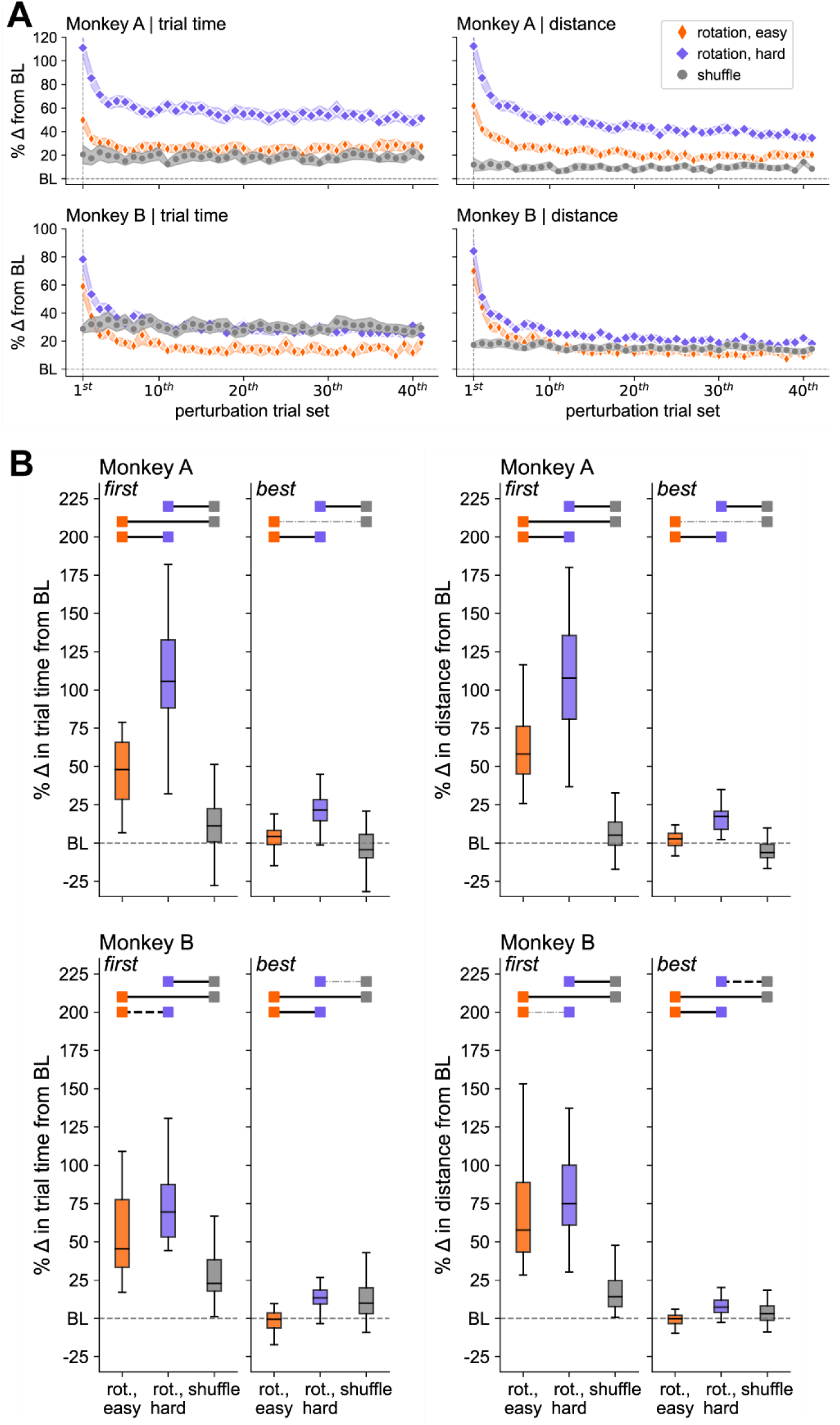
| Behavior under a shuffle perturbation does not exhibit a classical adaptation profile. In contrast to behavior under the rotation perturbation that exhibits a rapid exponential decay, behavior (over sets of 8 non-overlapping sets of perturbation trials) under the shuffle perturbation is essentially flat – indicating a lack of classical exponential adaptation. Solid black bars indicate significant differences, dashed black lines indicate trending differences (0.05<p<0.1) and grey dashed lines indicate no significant differences (one-way ANOVA, post-hoc pairwise comparisons (Tukey-Kramer adjustment)).

We then examined the behavioral performance during two sets of 8 perturbation trials: performance during the initial perturbation trial set and performance during the first the trial set in which performance was the best (i.e., closest to baseline (BL) or below). The rotation perturbation caused a larger performance deficit in behavior in comparison to the shuffle perturbation (**Figure 3.B**). Interestingly, under the shuffle perturbation, for Monkey A, 13/43 sessions started with initial performances better than baseline.

### Neurons exhibit a broad range of tuning changes in response to perturbation

To counteract the perturbation, subjects needed to alter the neural activity of their BCI neurons to restore performance. To quantify the neural responses, we assessed directional tuning changes of individual neurons pre– and post-perturbation (*see Methods – Tuning parameter estimation*). We found that, regardless of presence of classical adaptation, neurons exhibited a broad range of tuning changes when adapting to a perturbation.

Under the rotation perturbation, the average preferred direction change of neurons was proportional to the magnitude of the applied rotation and in the direction of the applied rotation (**Figure 4**). However, the session-average change was always less than the applied rotation. When examining changes in preferred direction of individual neurons, we observed a range of changes from no changes to changes in the opposite direction of the applied rotation condition to changes greater than the applied rotation (**Figure 5A,C**).

**Figure 4.**
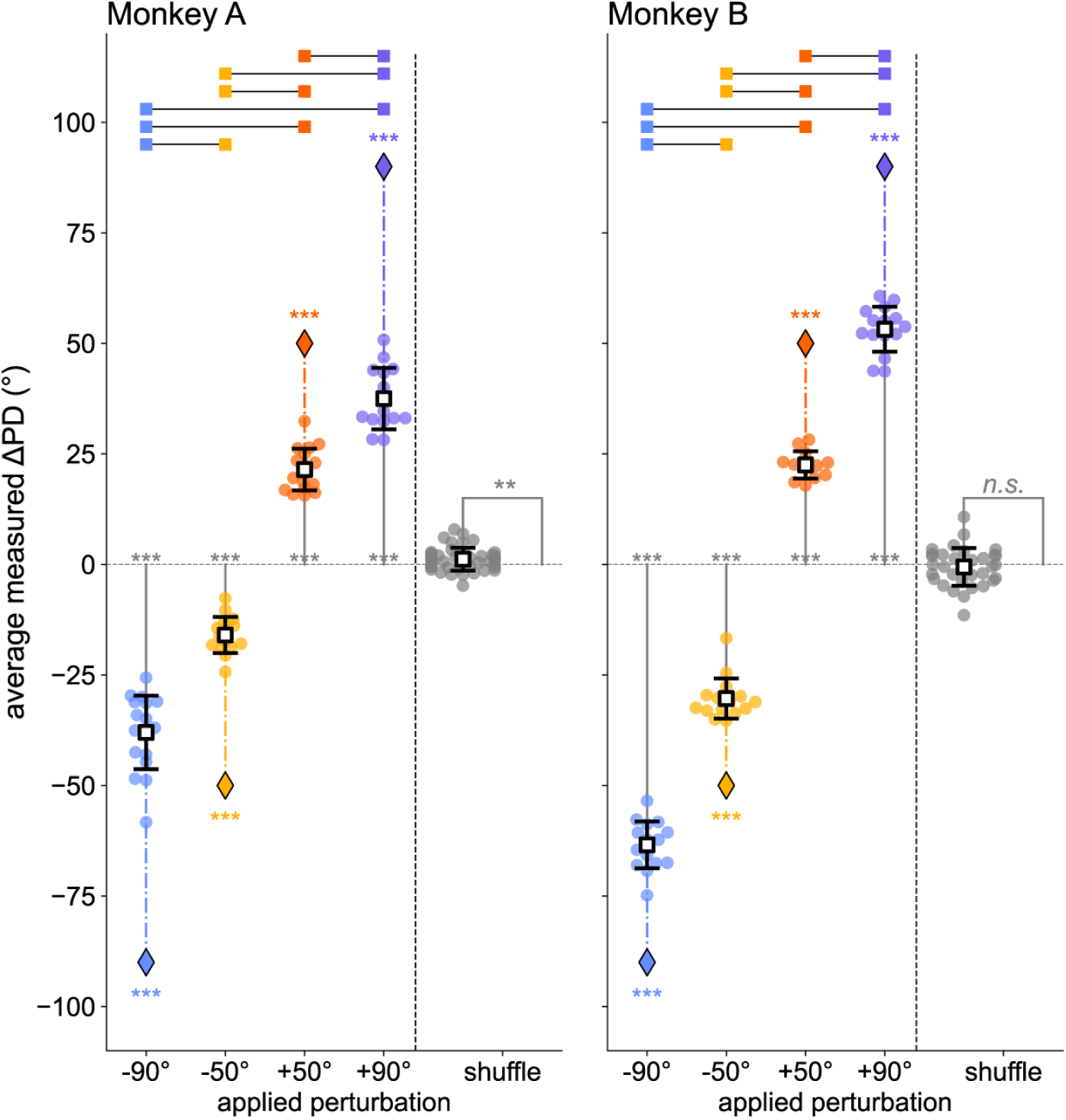
| Session-averaged changes in preferred direction under different decoder perturbations. Under the rotation perturbation, the average change in preferred direction is significantly greater than 0° and in the direction of the applied perturbation but less than the applied rotation amount. Under the shuffle perturbation, the average change in preferred direction may or may not be significantly different from 0° (depending on subject). However, the average change was small ( Monkey A: –4.8° to 7.8°; Monkey B: –11.5° to 10.7°). Each scatter point represents a session. See **Table 1** for number of sessions.

**Figure 5.**
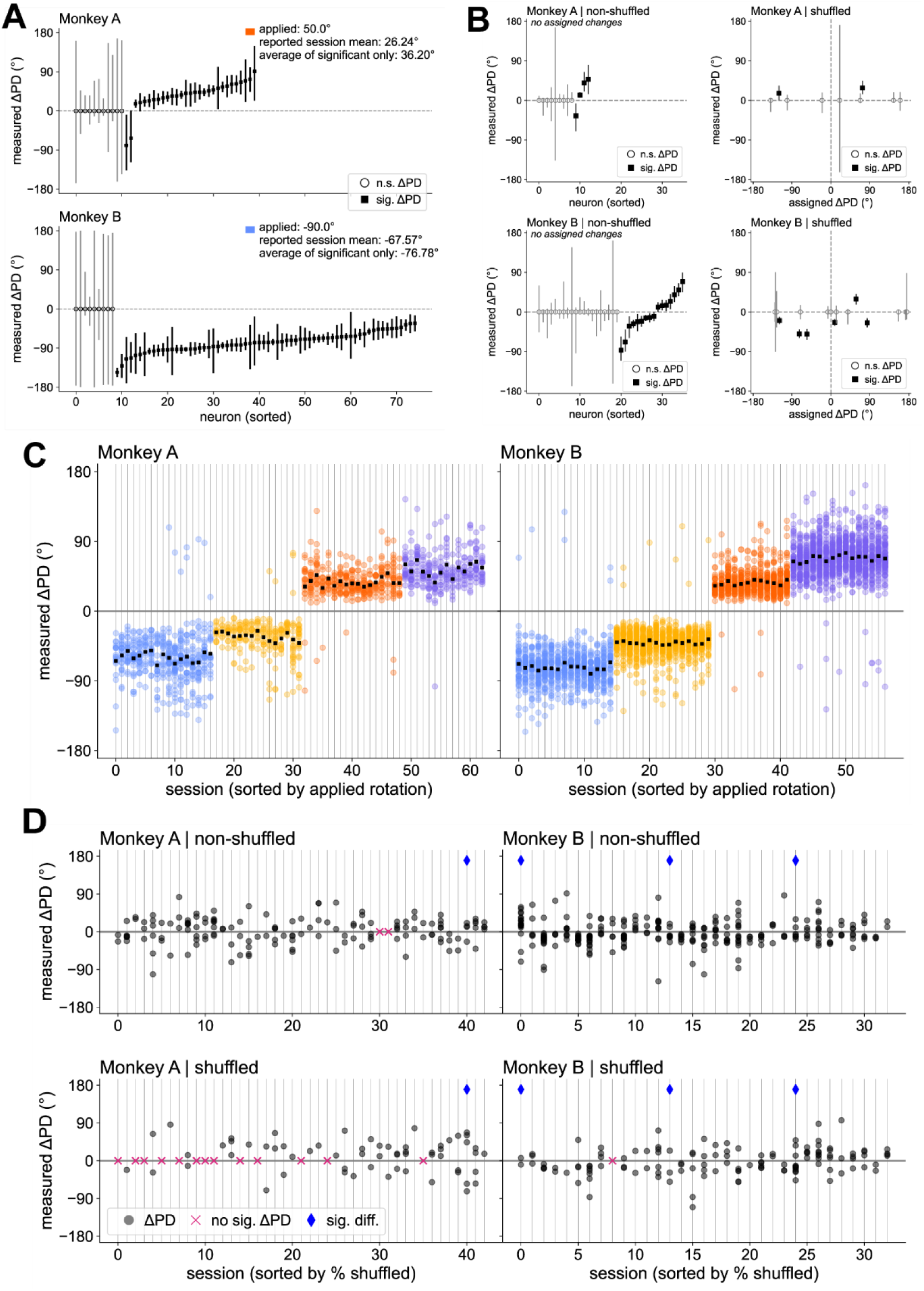
| Individual neurons display a range of preferred direction tuning changes. (**A**) Tuning changes of BCI neurons for a sample rotation perturbation session from each Monkey. Results presented as median measured changes in preferred direction and 95% confidence interval. (**B**) Similar to (A) for sample shuffle perturbation sessions. (**C**) Summary (rotation perturbation sessions) of all significant measured preferred direction changes per session, sorted by applied rotation angle. Black square represents population mean of significant changes. (**D**) Summary (shuffled perturbation sessions) of all significant measured preferred direction changes per session, sorted by percentage of shuffled neurons (Note that the number on the x-axis indicates session number – not percentage shuffled). Grey scatter points (circles) represent measured change in preferred direction. Magenta scatter points (x) represent sessions with no significant changes for non-shuffled or shuffled. Blue scatter points (diamond) represent sessions in which the average absolute value of significant preferred direction changes was significantly different between shuffled and non-shuffled neurons (Mann-Whitney U-test, p<0.05).

**Table 1.** | Number of completed session across perturbation types & average number of trials completed per block. “-“ indicates counterclockwise rotation and “+” indicates clockwise rotation. Except for **Figure 4** (to demonstrate that BCI neurons change preferred direction in the direction of applied rotation), +/− sessions were combined due to lack of statistical support for separating the conditions based on our previous behavioral analysis^24^. Trials beyond the baseline (BL) block and perturbation (PE) block were not used in the analyses presented here. They are reported to demonstrate that subjects were engaged in the task well beyond the analyzed trials. BL: baseline block, PE: perturbation block; non-EC: non-error clamp trials (The baseline block included “error clamp” (EC) trials in which the cursor error feedback was projected (“clamped”) onto a straight-line path to the target. These trials, which occurred once every eight trials, were excluded from analyses.) *Note: the baseline trial sets used to fit tuning curves were used to fit FA models and compute the two baseline metrics (%sv, %contribution). Even though subjects completed more trials under the rotation perturbations, the number of samples impacts both tuning estimates and variance estimates. Therefore, we used a constant number of trials across block and perturbation types to control for this*.

|  | Monkey A | Monkey B |
| --- | --- | --- |
| # sessions: rotation, easy (-/+50°) | 15 /17 (=32) | 15/12 (=27) |
| # sessions: rotation, hard (-/+90°) | 17/14 (=31) | 15/15 (=30) |
| # sessions: shuffle | 43 | 33 |
| # trials, BL: rotation | 328 non-EC (42 trial sets) | 328 non-EC (42 trial sets) |
| # trials, BL: shuffle | 168 non-EC (21 trial sets) | 168 non-EC (21 trial sets) |
| # trials, PE: rotation | 384 non-EC (48 trial sets) | 384 non-EC (48 trial sets) |
| # trials, PE: shuffle | 384 non-EC (48 trial sets) | 384 non-EC (48 trial sets) |
| # trials, beyond BL + PE: rotation (easy only) | 241.44 ± 136.42 | 235.87 ± 109.69 |
| # trials, beyond BL + PE: rotation (hard only) | 130.29 ± 132.26 | 237.27 ± 104.08 |
| # trials, beyond BL + PE: shuffle | 271.40 ± 183.51 | 404.73 ± 171.11 |

Unlike the rotation perturbation, the shuffled decoder effectively assigned a new preferred direction for a subset of neurons only (*Methods – Selecting neurons to shuffle*). Even though how much a single neuron should adapt its preferred direction is not obvious based on low-dimensional visual feedback, shuffled and non-shuffled neurons still significantly change their preferred direction in response to the shuffle perturbation. On average, the BCI population changed its preferred direction within a small range of degrees (**Figure 4**). Like neurons under the rotation perturbation, neurons under the shuffle perturbation exhibited a wide range of preferred direction changes (**Figure 5B,D**). In each session, we observed significant changes in preferred direction in non-shuffled and/or shuffled neurons. In 28 out of 44 sessions for Monkey A and 32 out of 33 sessions for Monkey B, we observed significant changes for both shuffled and non-shuffled neurons. Of these sessions only 1 (of 28) session for Monkey A demonstrated significant differences between shuffled and non-shuffled neurons; the absolute preferred direction changes were greater for shuffled neurons. Additionally, only 3 (of 32) sessions for Monkey B demonstrated differences between shuffled and non-shuffled neurons (n=2, |ΔPD| shuffled > |ΔPD| non-shuffled; n=1, |ΔPD| non-shuffled > |ΔPD| shuffled (session 0)). In the majority of these sessions, however, the absolute amount of change was not significantly different between the non-shuffled and shuffled neurons.

Examining the relationship between changes in assigned preferred direction versus the measured preferred direction changes revealed two key findings in the shuffled perturbation dataset. First, the non-shuffled neurons changed their preferred direction much more than expected. Specifically, the assigned preferred direction for non-shuffled neurons did not change under the perturbation, but we still measured a range of significant preferred direction changes (**Figure 6A**). Second, the shuffled neurons typically changed their preferred direction much less than expected (i.e., significantly different amounts than the assigned change; **Figure 6B**). In fact, we observed no significant relationship between the assigned and measured preferred direction change (p >> 0.05, Pearson’s correlation coefficient). Qualitatively, the majority of individual neuron preferred direction changes appear to be within ±45°, regardless of changes in assigned preferred direction. Given that we observed a wide range of preferred direction changes under the rotation perturbation, we would expect that individual neurons should be able to change their preferred direction in proportion to the assigned change. We do not expect that the neural limit in preferred direction change resides at the level of a single neuron. Rather, constrained changes likely reflect population-level constraints; neurons coordinate activity to counteract the perturbation and restore behavior with only a low-dimensional representation of the neural activity (i.e., which neurons are producing the error is unclear).

**Figure 6.**
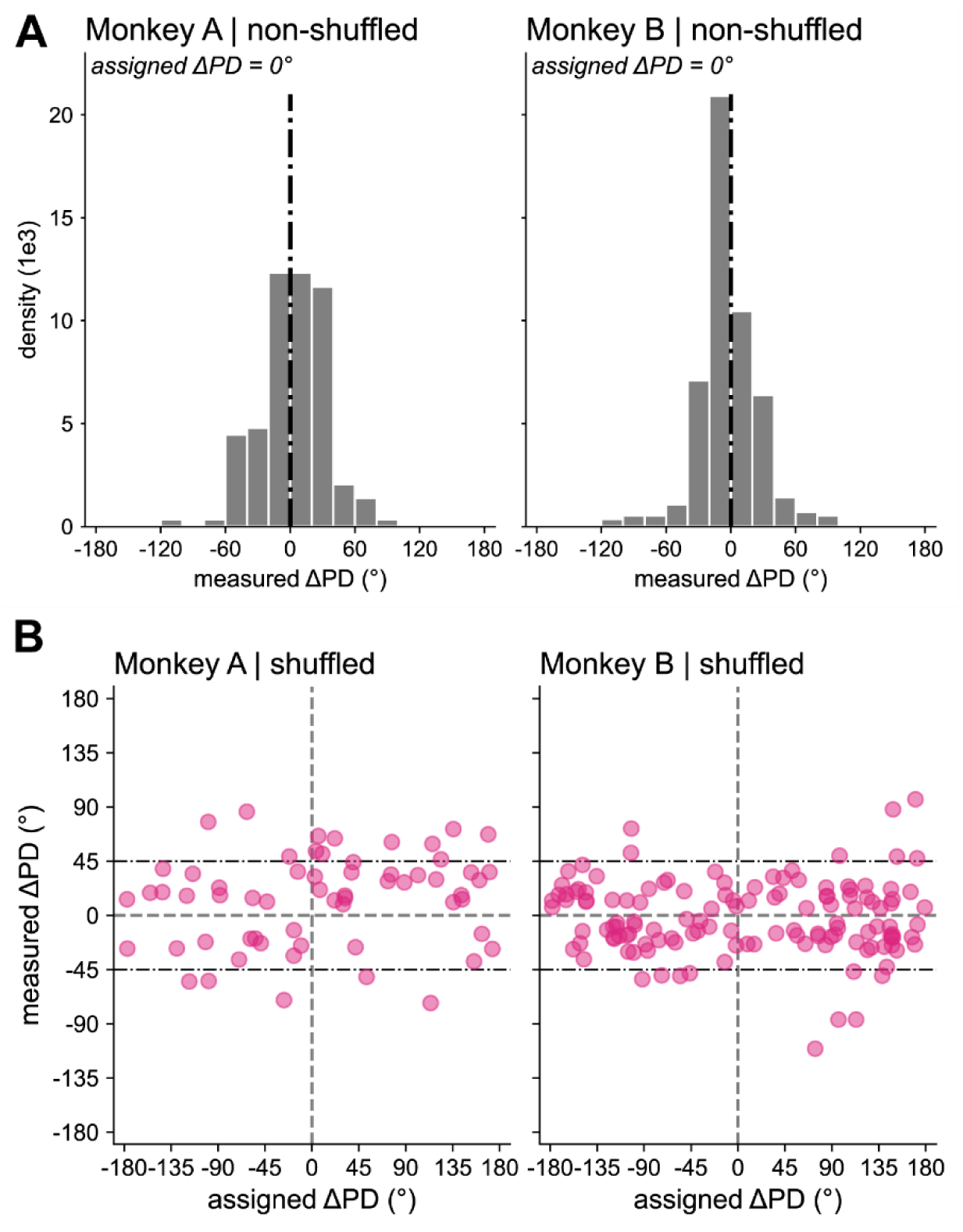
| Significant measured preferred direction changes in (A) shuffled and (B) non-shuffled neurons across all shuffle perturbation sessions. (**B**) We observed no significant correlation between assigned and measured changes in preferred direction changes in shuffled neurons (Monkey A, n=61, r=0.17, 0.18; Monkey B, n=130, r=-0.04, p=0.64; Pearson’s correlation coefficient).

Of the three perturbation types (easy rotation, hard rotation, shuffle), we observed the highest percentage of significant changes under the hard rotation condition (**Figure 7A**). One factor that impacts the ability to measure small changes is the modulation depth^38^. To ensure that our results were not reflective of measuring poorly-tuned neurons (i.e., low modulation depth) we analyzed modulation depth during the baseline block, during the perturbation block, and any changes between the two blocks. The session-averaged modulation depth was significantly greater under the shuffle condition for both baseline and perturbation tuning models and was not significantly different between rotation types (**Figure 7B**). We found that neurons that did not change their preferred direction also did not change their modulation depth significantly (**Figure 7C,D**). This was also true for neurons with significant changes in preferred direction under the easy rotation condition. In contrast, neurons with significant changes in preferred direction under the hard rotation condition significantly increased their modulation depth. (Given the large sample size, we would like to note that the effect size of this comparison is small (Monkey A, d=0.21; Monkey B, d=0.25; Cohen’s d)).

**Figure 7.**
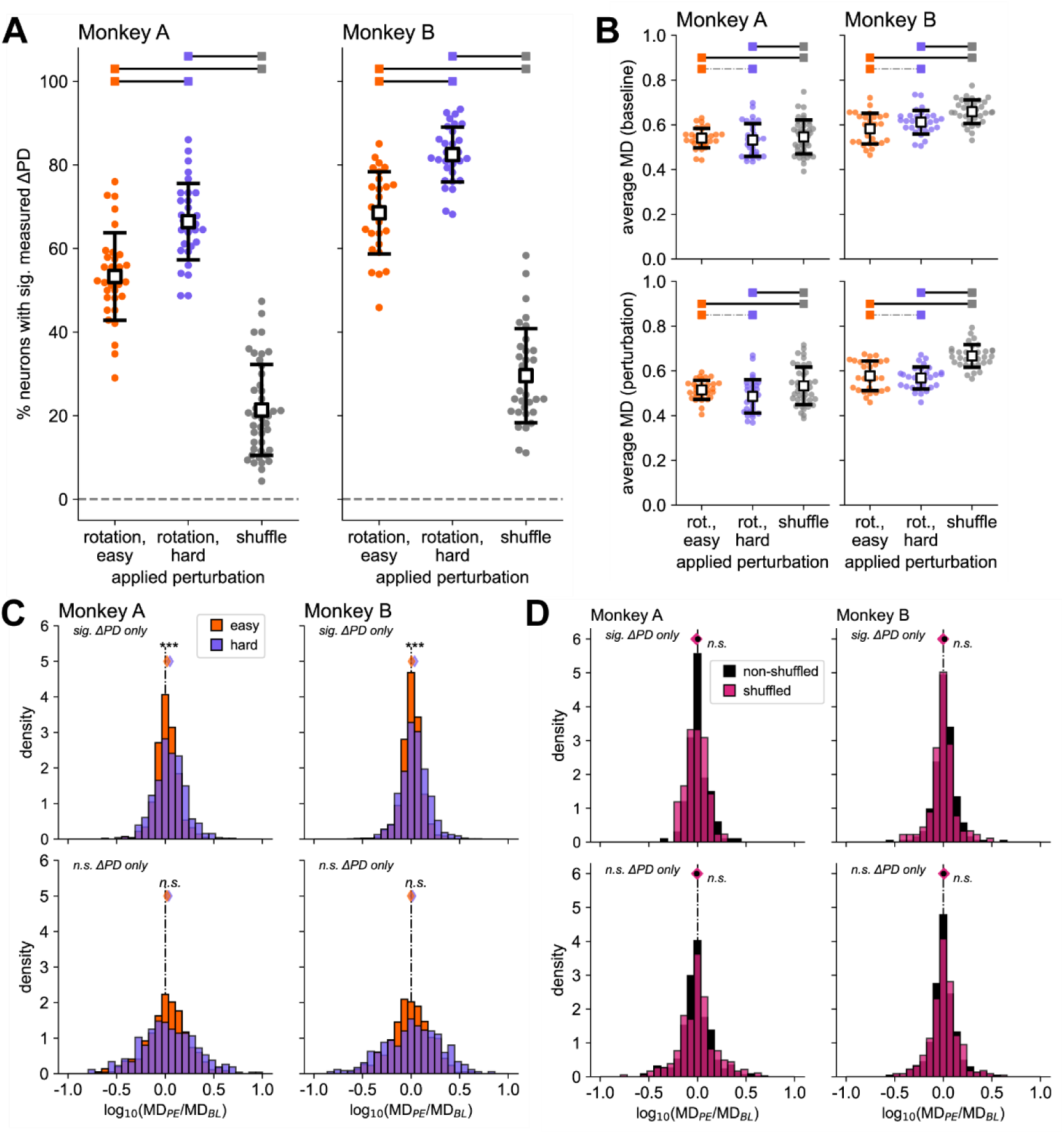
| The difference in the number of neurons with significant tuning changes across perturbation types cannot be explained by poor tuning. (A) Percentage of BCI neurons with significant changes in tuning. Each scatter point represents a session. (B) Average measured modulation depth for both the baseline and perturbation tuning models. Means (± standard deviation) are represented by a black-outlined white square. Black lines indicate significant difference and grey dot-dash lines indicate no significant difference. (C) Under the rotation perturbation, neurons with non-significant (*n.s.*) preferred direction changes also exhibited no significant changes in modulation depth. Changes in modulation depth were only significant (*sig.*) for neurons under the hard rotation condition that also had significant preferred direction changes. (D) Under the shuffle perturbation, we observed no significant changes in modulation depth.

**Figure 7.**
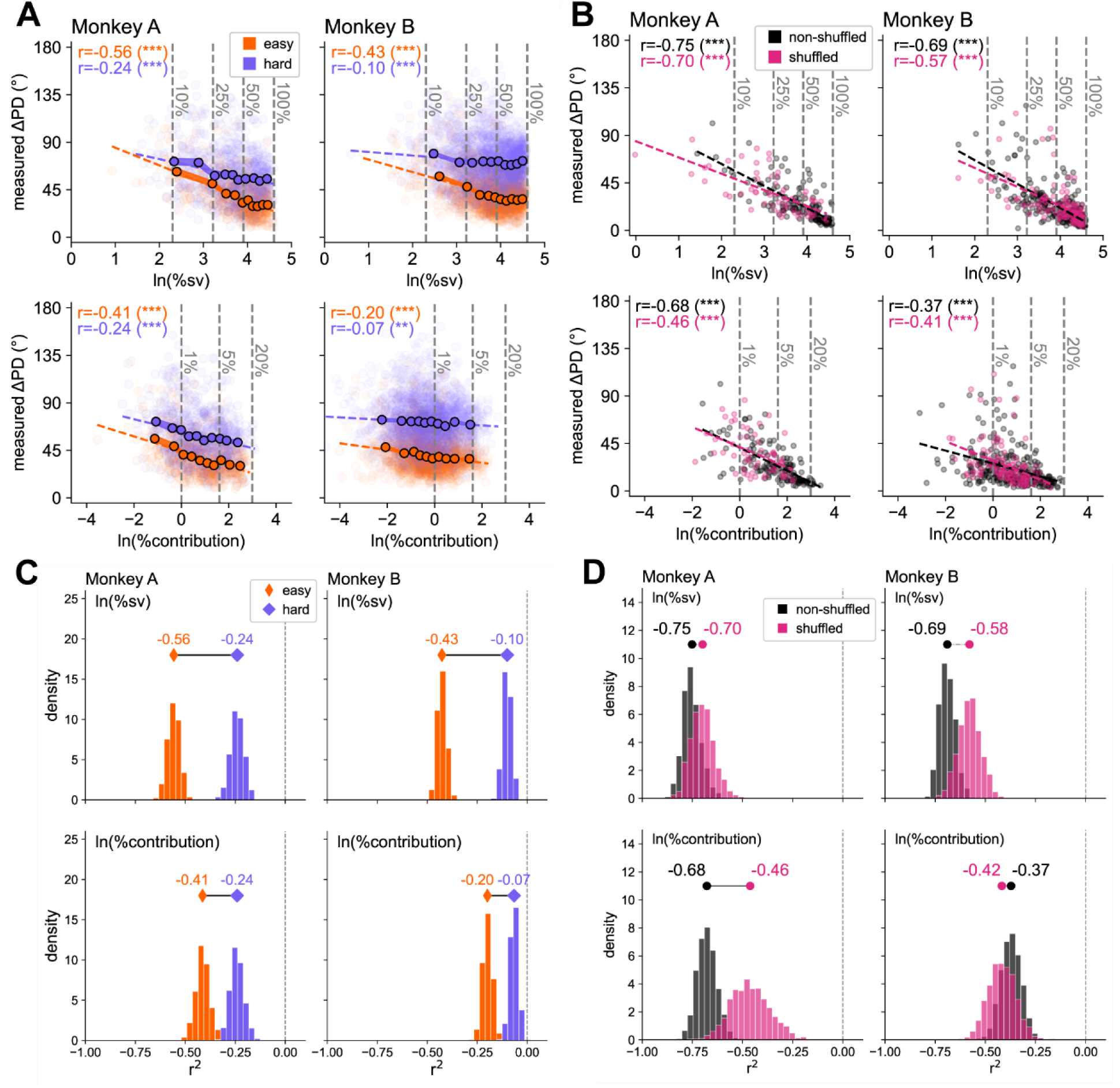
| Neural baseline metrics are highly correlated with the change in preferred direction. (**A**) Under the rotation perturbation, neurons generally changed their preferred direction more if they had low %sv and contribute little to the decoder output. (**B**) Similarly, for non-shuffled and shuffled neurons. (**C**) Bootstrapping revealed that these metrics explained more of the variance observed in preferred direction changes under the easy condition. (**D**) The correlative relationship between each metric and the amount of change in preferred direction was only significantly different for Monkey A, decoder contribution (ln: natural log, %sv: percent shared variance, %contribution). *For the precise number of neurons used in these analyses, see* Table 2.

**Table 2.** | Number of neurons by perturbation type. Average (±standard deviation) number of neurons per session included (or excluded based on < 1Hz firing rate) for easy rotation session, hard rotation session, and shuffle session. sig. measured ΔPD (total): total number of included neurons with significant measured changes in preferred direction within condition (total number of neurons within condition).

|  | **perturbation type** | **Monkey A** | **Monkey B** |
| --- | --- | --- | --- |
| mean # neurons, included | rotation, easy | 35.38 ± 6.98 | 95.15 ± 12.62 |
| mean # neurons, *excluded* | rotation, easy | 0.91 ± 1.07 | 2.44 ± 1.87 |
| total # neurons, included | rotation, easy | 1132 | 2569 |
| total # neurons, with sig. measured ΔPD | rotation, easy | 599 | 1759 |
| mean # neurons, included | rotation, hard | 36.39 ± 6.96 | 89.90 ± 25.79 |
| mean # neurons, *excluded* | rotation, hard | 1.26 ± 0.88 | 2.03 ± 1.64 |
| total # neurons, included | rotation, hard | 1128 | 2697 |
| total # neurons, with sig. measured ΔPD | rotation, hard | 745 | 2189 |
| mean # neurons, included | shuffle | 22.70 ± 4.37 | 42.12 ± 10.52 |
| mean # neurons, *excluded* | shuffle | 0.98 ± 0.88 | 0.91 ± 1.46 |
| total # neurons, included | shuffle, shuffled neurons | 325 | 450 |
| total # neurons, with sig. measured ΔPD | shuffle, shuffled neurons | 61 | 130 |
| total # neurons, included | shuffle, non-shuffled neurons | 651 | 950 |
| total # neurons, with sig. measured ΔPD | shuffle, non-shuffled neurons | 146 | 282 |

Regardless of perturbation type or the observation of expected behavioral adaptation, neurons adapted to external changes when presented with low-dimensional feedback. Furthermore, neurons, on average, primarily change the direction in which they fire the most (i.e. change preferred direction) but not necessarily the rate of firing. Interestingly, even when there was no mismatch between the decoder and the neural activity responsible for behavior (i.e., non-shuffled neurons), we observed significant tuning changes. We next investigated the factors that constrain (or support) the ability of neurons to adapt their activity appropriately.

### Baseline characteristics predict amount of preferred direction changes

If each neuron acted independently to counteract the perturbation, then the optimal solution for adaptation would be for each neuron to alter its preferred direction by the applied rotation angle to restore behavioral performance. However, we know that the neurons are embedded in a highly coordinated network, likely facilitating individual neurons to adapt their preferred directions in a way that reflects the collective dynamics rather than by the exact amount of the applied perturbation. We investigated how two measures of baseline neural activity impacted the “mismatch” between the applied and measured preferred direction changes. The first metric, percent shared variance (%sv), quantified how coordinated a neuron’s activity was with the BCI population activity (*see Methods – Computation of % shared variance (%sv) through factor analysis*). The second metric, percent contribution (%contribution; *see Methods – Computation of %contribution to behavioral output*) acted as a measure of a neuron’s task-relevancy. Specifically, we computed the average population neural command magnitude and then determine how much each neuron’s weighted spiking activity contributed to the neural command magnitude (computed within session).

Initial examination of the results revealed similar relationships between the two metrics and amount of measured preferred direction change (**Figure 7**). Interestingly, we observed the lowest amounts of preferred direction changes in neurons with the high levels of population co-variability – regardless of the applied perturbation. Similarly, neurons with larger contributions to the behavioral output also changed their preferred direction the least. Generally, these two metrics were positively correlated. (Monkey A, r_easy_=0.67, r_hard_=0.66, r_shuffled_=0.51, r_non-shuffled_=0.76; Monkey B, r_easy_=0.57, r_hard_=0.57, r_shuffled_=0.62, r_non-shuffled_=0.61; Pearson’s correlation coefficient, p<<<0.05). Therefore, neurons with the most population-level and behavioral-output influence, still changed their preferred direction, but changed preferred direction the least. Under the shuffle perturbation, for non-shuffled neurons, minimal changes are desired (from a single neuron perspective), as we did not apply a perturbation directly to these neurons. Conversely, by nature of connections with these neurons, shuffled neurons were increasingly limited in the ability to independently change their preferred direction.

### Mismatch of preferred direction changes predicts behavioral improvements

To demonstrate that measured changes in individual neurons reflect behavior, we determined how mismatch between assigned and measured preferred direction changes related to behavioral changes under both perturbation types. For both the rotation and shuffle perturbations, lower amounts of mismatch predicted better behavioral outcomes under the perturbation (**Figure 8**). Specifically, lower levels of mismatch in preferred direction changes corresponded to faster trial times and shorter distances. Importantly, the mismatch metric can be used to predict how well the subject can perform within a session under all perturbation types.

**Figure 8.**
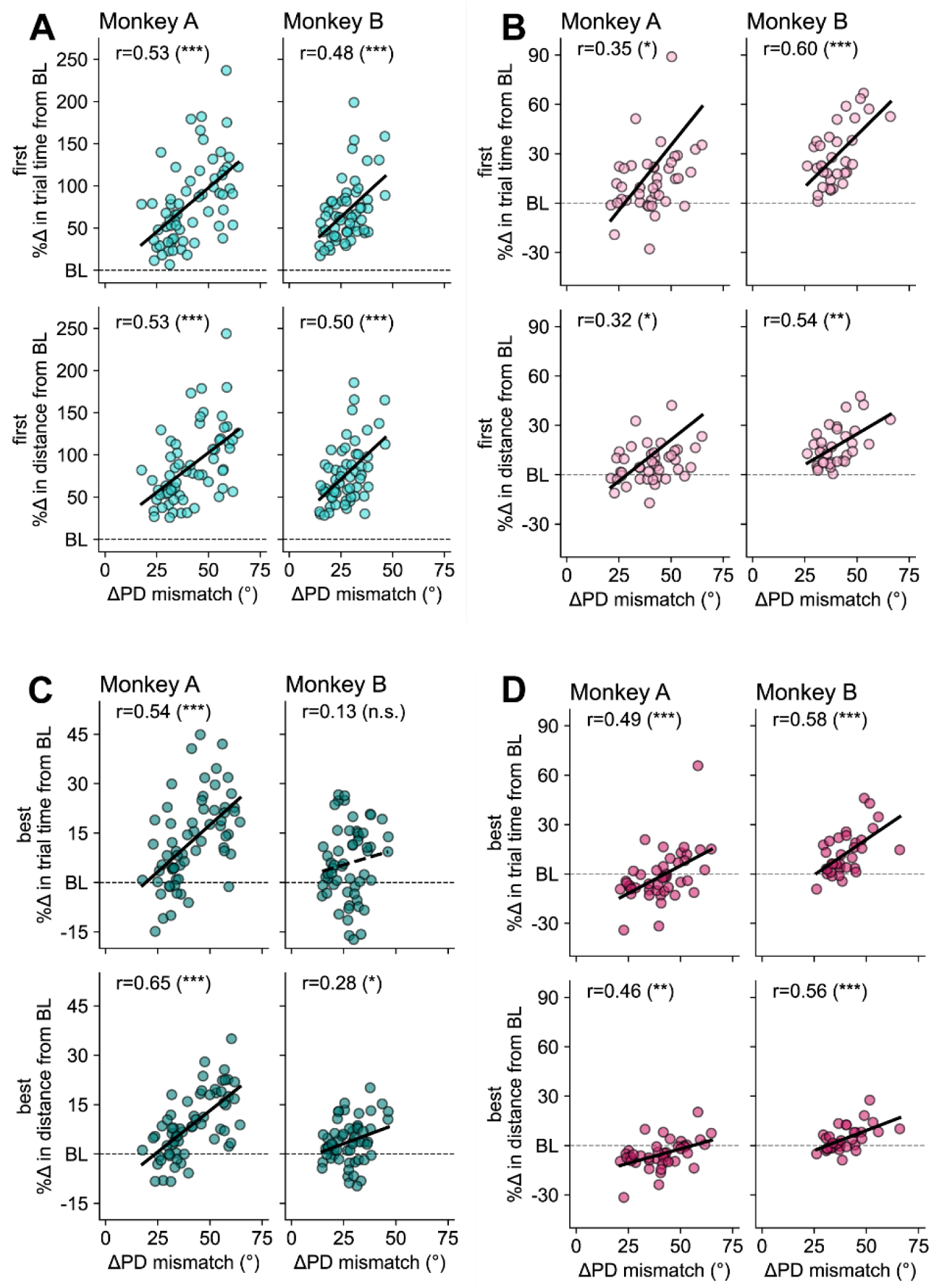
| Accuracy of preferred direction changes predicts behavioral improvements. Performance during the *initial* perturbation trial set under both rotation (**A**) and shuffle (**B**) perturbations and the *best* performance under both the rotation (**C**) and shuffle (**D**) perturbations are positively correlated with the session-averaged preferred direction change mismatch of individual neurons. Each scatter point represents a session. One session (Monkey A, shuffle perturbation) was removed from visualization (%Δ >> 100); however, the line and statistics presented on the plot include this session. r: Pearson’s correlation coefficient.

In line with our tuition that neurons work together to counteract the perturbation, we found that when we computed the session-average change in preferred direction and then computed mismatch from the global rotation signal (i.e., ± 50 or 90°), we obtained similar results as in **Figure 8**. Conversely, under the shuffle perturbation, when we computed the session-average change in preferred direction and then computed mismatch for the global shuffled signal (computed as the average assigned preferred direction change over shuffled and non-shuffled neurons), the results did not hold. While we concede that computing a simple mean is not representative of the low-dimensional neurofeedback signal, this does highlight that the activity of individual neurons within a population-context are important to study to ensure careful design of neurofeedback signals and protocols that do not disparately impact sub-populations within the neurofeedback population.

## DISCUSSION

Here, we investigated how baseline characteristics influenced the ability of single neurons to adapt under different types of perturbation and, in turn, how this ability resulted in behavioral changes. Specifically, we explored the relationship of baseline metrics –%sv and %contribution (to behavioral output)– on the amount of preferred direction changes. We discovered a remarkably similar relationship across perturbation types in which increased values of the baseline metrics predicted smaller changes. Overall, better accuracy of tuning changes was predictive of better performance under the perturbation and better overall recovery in performance.

We first observed that average behavior under the shuffle perturbation did not follow an exponential trend that is typical of the short-term adaptation observed like under the rotation perturbation, in other non-human primate studies^10,11,26^, and in human studies^27–37^. Given the nature of the shuffle perturbation, we expected that the dimensions of neural activity would be disrupted in a non-uniform fashion. Similar perturbations, that completely violate the underlying biological neural network structure, typically require multiple sessions of practice to achieve adequate control^12,13,18^. However, we demonstrate here that under more subtle violations, monkeys can perform sufficiently, although with limited amounts of improvements (**Figure 2**, **Figure 3**).

Even in the absence of classical temporal adaptation, we observed a wide range of preferred direction changes under the shuffle perturbation (**Figure 5**, **Figure 6**). Overall, significant changes in preferred direct were not selective to perturbed neurons, potentially due to the inability of neurons to perfectly resolve low-dimensional feedback to the specific perturbed neurons (“credit-assignment problem”^39^). In fact, we did not observe significant evidence that shuffled versus non-shuffled neurons changed more. This is somewhat in contrast with previous studies that demonstrate that perturbed neurons change their preferred direction more than non-perturbed neurons^10,11^ and that neurons directly mapped to BCI behavior modulated their activity more than surrounding neurons not used for BCI control^9^. Even though we were able to effectively measure small changes in preferred direction our results are likely impacted by the small number of neurons exhibiting significant preferred direction changes. Still, these results still support the idea that, just as in natural motor behaviors, BCI neurons are constrained by their underlying network, which in turns impacts the amount of possible behavioral changes. Given the growing body of evidence that successful BCI control and adaptation are constrained by neural activity from regions not directly responsible for BCI behavioral output, future investigation of the constraints caused by other cortical areas (e.g., sensory cortices) and sub-cortical areas^40^ (e.g., striatum) is warranted.

As many as half of participants in neurofeedback training are unable to modulate their neural activity effectively (“non-learners”)^6,41^. Furthermore, the ability to adapt neural activity does not necessarily result in desired behavioral outcomes^42^. Our new line of evidence contributes to increased understanding of the constraints of single-neuron modulation by identifying reliable population-level predictors of an individual neuron’s ability to adapt appropriately and to produce desired behavioral outcomes within a single day. Furthermore, we have previously demonstrated^43^ that techniques, similar to the ones presented here, can be implemented on data from different neural recording modalities (i.e., functional magnetic resonance imaging versus implanted microelectrode arrays) and on different species (i.e., humas, non-human primates) and decode meaningful task-relevant information. Therefore, we are optimistic that these methods and findings can be translated beyond the realm of implantable BCIs in non-human primates.

## METHODS

### Code availability

All experiments and analyses were run using Python3 (version 3.9.7). Analysis and plotting scripts are available upon request. Specific Pythonic libraries and functions are provided in the sections below. Original experimental code (BMI3D library) can be found here: https://github.com/carmenalab/brain-python-interface, and customized experimental code can be found here: https://github.com/santacruzlab/bmi_python.

### Experimental model: Rhesus macaques

Two group-housed, healthy male rhesus macaques (*Macaca mulatta*) were used to perform all studies (Monkey A: age 5, ∼9.5kg; Monkey B: age 5, ∼9.5kg). All rhesus macaque procedures were approved by the Institutional Animal Care and Use Committee (IACUC) of The University of Texas at Austin, a fully AAALAC-accredited institution. The experiments were conducted in accordance with the *Guide for the Care and Use of Laboratory Animals*, Public Health Service Policy, and the Animal Welfare Act and Regulations. Full surgical and training records can be found in our previous work^24^. Briefly, monkeys were implanted with chronic, Utah-style microelectrode arrays (Innovative Neurophysiology Inc.; Durham, NC) in the left primary motor (M1) / pre-motor (PMd) forelimb cortical areas. From these arrays, we recorded discrete “spike” events (i.e.,) from pre-sorted neurons (single or multi-unit) using the Grapevine Neural Interface Processor (Ripple Neuro; Salt Lake City, UT).

### Behavioral metrics

To quantify behavioral performance, we analyzed how two common performance metrics changed as a result of a decoder perturbation that was introduced at the start of the perturbation block. We defined the trial duration (“trial time”) as the amount of time elapsed from the onset of the appearance of the peripheral target (end of “hold center”, **Figure 1B**) to the start of the peripheral target hold period. Additionally, we defined cursor path length (“distance”) as the sum of the straight-line distances between each cursor position over the same window in which trial time was computed.

We measured baseline performance by averaging each metric over all baseline block trials. We examined performance during the perturbation block by computing average performance metrics within each perturbation trial set. We defined a “trial set” as 8 consecutive trials (i.e., 1 per target location). Trial sets did not overlap. Additionally, for each trial set, we computed the percentage change from the average baseline metric to allow for fair comparison across sessions. We present behavioral results as percentage change in trial time (or distance) relative to the mean baseline trial time (or distance). Increased trial times (or distances) relative to baseline indicated worse performance, and values closer to or below baseline (with baseline represented by “0”, by definition) indicated performance improvements.

### Kalman filter decoder

The Kalman Filter (KF) is commonly used to linearly map spike vector data of neurons used for BCI control (“BCI neurons”) to computer cursor velocity^12,44,45^. Once the KF has been trained, the resultant model can be expressed as a steady-state, linear, time-invariant form as in **Equation 1**,

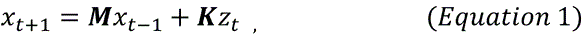

where **x** is a vector containing the horizonal and vertical components of cursor movement; **M** is a matrix of constants that weights a portion of the previous state to the current timestep, *t*; and **K** is the Kalman gain matrix that assigns an x– and y-velocity weight to each neuron’s spike counts activity (*Z*_*t*_). Cursor position was updated every 10Hz based on the newly estimated velocity.

We trained a new KF decoder at the start of each experimental session (day) using data collected from monkeys passively observing a cursor completed trials (“visual feedback task”, VFB; 6 trials x 9 targets; 3.5 minutes). To further tune the decoder, we implemented closed-loop decoder adaptation (CLDA) prior to starting the main perturbation task^21,24^. CLDA was not applied during the main task.

### Decoder perturbation types

In this body of work, we implemented different types of perturbations to study neural adaptation underlying behavioral adaptation. We implemented a rotation perturbation in which we tested an easy rotation perturbation (defined by a rotation angle of 50°) and hard rotation perturbation (defined by a rotation angle of 90°)^24^. To implement a rotation perturbation, we applied a rotation matrix (***R***(***θ***)) that altered the gain for each BCI neuron (**Equation 2**):

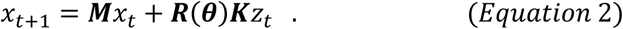

This altered the horizontal and vertical gain for each BCI neuron to change. Prior to adaptation of neural activity, the rotation perturbation initially caused the cursor position to appear rotated to the intended direction.

We also presented results from a shuffle perturbation task. Within a subset of BCI neurons, we randomly permuted the rows of **K** (i.e., the assigned weights for the x– and y-components of velocity). The effects of the shuffle perturbation on cursor position varied by session. Notably, we chose to shuffle a subset of BCI neurons because we were primarily interested in studying changes on the short timescale of adaptation (within-day) rather than the long-term learning (across days) that would be required to learn proficient control of an entirely novel (i.e., completed shuffled) decoder ^13,18^. For a given shuffle perturbation session, we selected approximately 18% to 50% of neurons to undergo shuffling. To select the neurons to be shuffled, we first computed average firing rates and preferred directions from the VFB data. Neurons below the 10^th^ percentile and above the 90^th^ percentile were excluded from shuffling. The remaining neurons were sampled pseudo-randomly to ensure that the PD of shuffled neurons were not clustered in a single direction and were, instead, distributed as equally as possible in all directions. At the onset of the perturbation block, the Kalman gain for shuffled neurons were reassigned within the group of shuffled neurons.

### Tuning parameter estimation

The directional tuning of motor cortical neurons is commonly and sufficiently modeled by a cosine function^38,46–50^. By fitting a cosine curve to firing rates computed for multiple movement directions, we can derive two parameters. The preferred direction (PD) is the movement direction, in degrees, corresponding to the maximum firing rate. The modulation depth (MD) is the amplitude of the curve divided by the baseline firing rate and represents how selective the neuron is to its PD.

For our task, we computed firing rates for each neuron for each of 8 target locations, which represent intended movement directions. The tuning curve was fit to the mean firing rate for each target direction (θ) as in **Equation 3**^18,21,38,51^:

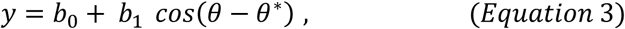

where the firing rate (y, spikes/second) is approximated by a mean firing rate across all movement directions (*b*_*o*_, spikes/second), a cosine modulation factor (*b*_1_) and the phase shift (θ^∗^), which is interpreted as the PD. We then computed the modulation depth as *b*_1_⁄*b*_*o*_ from **Equation 3**. In practice, we estimated tuning curve parameters by implementing least squares optimization through the *curve_fit* function from the SciPy statistics library (version 1.7.0).

To estimate the tuning parameters for each neuron, we fit cosine curves to bootstrapped samples of firing rates^38^. We first computed the neuron’s firing rate for each trial over an equal number of trials to each target direction (20 trials x 8 target directions = 160 firing rate estimates). The last 160 trials from each block were used. Using late trials avoids the period of rapid changes during early adaptation in perturbation. For each trial, we summed the neuron’s spike counts over a fixed window of 400ms and divided by the window length to compute firing rate. The start of this window was aligned to 200ms post the start of the movement cue. We excluded the first 200ms from analysis to account for the monkey’s processing delay and capture activity that is most reflective of the intended target direction. Once we computed the estimated firing rate for each trial, we implemented the bootstrapping procedure. Specifically, we randomly sampled, with replacement, the firing rate from 20 trials for a target location and then computed the mean. We repeated this process for each of the 8 target locations. Then, we fit a cosine curve to the 8 bootstrapped means. We repeated this process 1000 times to generate a distribution of PDs and MDs for each neuron’s activity under each block (1000 baseline estimations, 1000 perturbation estimations).

To estimate the changes in the PD from before to after the perturbation, we computed the smallest angular distances between a randomly-selected sample from the baseline bootstrap distribution and a randomly-selected sample from the perturbation bootstrap distribution. The result of repeating this process was a distribution of 1000 changes in PD. We defined changes in the clockwise direction as negative and changes in the counterclockwise direction as positive. To assess the significance of the change, we computed the 95% confidence interval (CI; significance level, α = 0.05). We considered a neuron’s change in PD as significant if the 95% CI did not include 0, and we reported the measured change in PD as the median of the bootstrapped distribution. For neurons with non-significant changes, we reported the change in PD as 0. Typically, the median value for these neurons were close to 0.

### Assigned tuning changes computation

When applying a decoder perturbation, we effectively “assigned” a new preferred direction to each BCI neuron by altering the values of **K**. Under the rotation perturbation, the rotation matrix effectively re-weighted the x– and y-components of Kalman gain for each neuron. Under the shuffle perturbation, we randomly permuted the rows of the **K** among a subset of neurons. Under both cases, this perturbation was applied once at the onset of the perturbation block and remained fixed throughout the perturbation block.

Under the rotation perturbation, by definition, the assigned PD change for each neuron was equal to the degree of the imposed rotation perturbation: |50°| (easy condition) or |90°| (hard condition). Under the shuffle perturbation, the assigned PD changes were zero for non-shuffled neurons and were unique for each shuffled neuron. To compute assigned PD changes for shuffled neurons, we first computed the assigned PD^21^ using the gain associated with each neuron under the baseline condition as in **Equation 4** and then under the perturbation condition as in **Equation 5**:

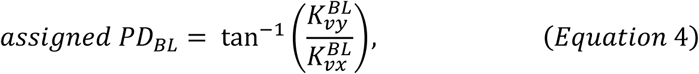

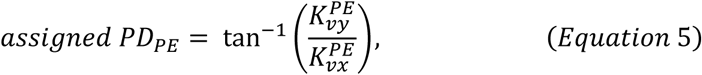

where **K** represents the Kalman gain, the superscript represents the block (*BL*: baseline, *PE*: perturbation), and the subscript represents the x– or y-component of velocity.

We resolved the PD to the appropriate Cartesian quadrant based on the signs of the inputs. We then computed changes in PD by finding the smallest angular difference between the two angles. We defined clockwise rotations as negative and counterclockwise rotations as positive.

### Computation of % shared variance (%sv) through factor analysis

Recent technological advances that enable researchers to record from more neurons simultaneously have pushed neural variability analysis towards methods that can summarize (or reduce the dimensionality of) the activity of tens to thousands of neurons^52,53^. One popular dimensionality reduction method is principal component analysis (PCA). This method summarizes the joint activity by finding the dimension(s) of activity that explain the most variance in the population activity. Principal components are linear combinations of the data and make no distinction between unique and common sources of variability. A related dimensionality reduction method, factor analysis (FA), explicitly accounts for neural variability that is shared across neurons and neural variability that is unique to individual neurons. Additionally, factors describe unobserved (latent) dimensions of activity that impact the population rather than summarizing population activity. These factors can be thought of representing common inputs that impact the joint activity of neurons.

Factor analysis (FA) is a useful dimensionality reduction method for identifying how neurons within a measured population co-vary their activity while also measuring variability that is unique to individual neurons. We used FA to decompose neural population activity into mean, shared (correlated), and private (uncorrelated) components for each BCI neuron. We modeled population activity as in **Equation 6**^15,54,55^:

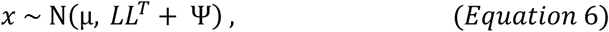

where *x* ∈ ℝ^*N*^ is the spike counts of *N* neurons, *μ* ∈ ℝ^*N*^ is a vector of mean spike counts for each neuron (*N*) in the population, ***L*** ∈ ℝ^*N*×*Q*^ represents the factor loading matrix of *Q* factors (where Q < N), and Ψ ∈ ℝ^*N*×*N*^ is the private variance matrix. To estimate the model parameters, we used the expectation-maximization (EM) algorithm ^54–56^.

We defined the shared variance and private variance as in **Equation 7**, and **Equation 8**, respectively:

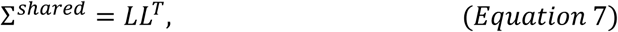

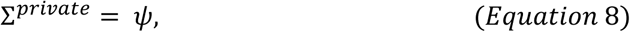

We fit one FA model per session on baseline block data. The dimensionality of the input was defined by the number of trials and the number of BCI neurons. Each entry in the matrix was the number of spikes observed within a fixed window of 400ms window time for one neuron. The same window and number of trials were used as in the tuning curve analysis.

To determine the number of factors used to fit each model, we performed repeated 10-fold cross-validation (CV) to determine the number of factors that maximized the average log-likelihood score of the model fit. We then chose the number of factors needed to explain 95% of the shared variance explained by the model that was fit using the number of factors identified through CV. This is a common practice to reduce overfitting, as the amount of shared variance explained by each additional factors tends to saturate below the value that maximizes the fit.^15,54,55^ For each neuron, *i*, we reported the percentage of variance that was explained by the factors over the total variance (percent shared variance, %sv) as in **Equation 9**:

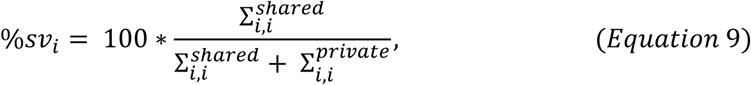

where *i* is the index of an individual neuron, and shared (**Equation 7**) and private variance (**Equation 8**) matrixes described above.

In practice, we used the *cross_val_score* function from the *scikit-learn* library (version 1.0.1) to perform CV, and the *FactorAnalysis* function from the *scikit-learn* library (version 1.0.1) to fit FA models.

### Computation of %contribution to behavioral output

For each session, we measured how each neuron contributed to the average total behavioral output under the baseline decoder. To compute the contribution each neuron had to behavior, we investigated the neural command (***K**Z*_*t*_; from **Equation 1**). We first determined the average spike count vector produced by neurons on each update of the decoder (i.e., every 100ms), and denoted this vector as *Z*. This average was computed over the same window and trials from the baseline block as used to calculate %sv and tuning changes. We then computed the average neural command for each neuron, *i*, for the x– and y-directions as in **Equation 10** and **Equation 11**, respectively:

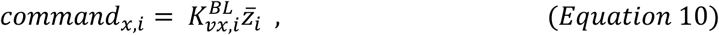

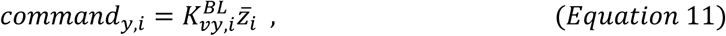

and then computed the magnitude of each neuron’s contribution as in **Equation 12**:

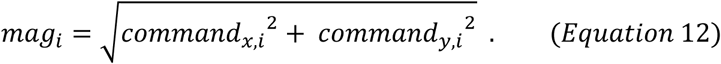

To compute the average total neural command magnitude, we summed the individual neuron magnitude contributions over all *N* BCI neurons as in **Equation 13**:

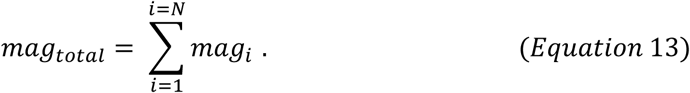

Finally, we computed and reported the contribution of each neuron as in **Equation 14**:

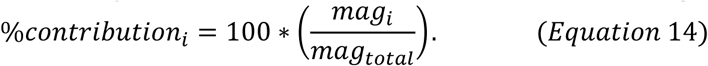

### Statistical methods

We performed all statistical and mathematical modeling in custom-written Python 3 scripts using the *SciPy* library (version 1.7.0). The statistical details for each figure can be found in the figure legends or the table corresponding to the figure. In general, we assumed a significance level of α = 0.05. Non-significant (*n.s.*) results were defined by p-values greater than or equal to 0.05. One star (*) indicated a p-value less than or equal to 0.05 but greater than 0.01. Two stars (**) indicated a p-value less than or equal to 0.01 but greater than 0.001. Three stars (***) indicated a p-value less than or equal to 0.001.

Prior to any two-sample t-tests for mean differences, we evaluated the ratio of sample variances using an F-test. If the F-statistic was less than or equal to three, sample variances were considered equal, and we used a Student’s t-test. Otherwise, we assumed sample variances were unequal, and we used Welch’s t-test. ANOVAs (two-way, one-way) were performed with post-hoc pairwise comparisons of means using a Tukey-Kramer adjustment.

In practice, we tested both positive and negative rotation conditions. However, we combined the positive and negative rotations based on the lack of statistical support for separating the conditions in both subjects based on behavior. We previously found that the main effects were limited to the trial sets (two-way ANOVAs^24^).

### Summaries of number of sessions, number of trials, and number of neurons by perturbation type

In this section, we present two tables that summarize the number of data points used for statistical modeling and testing. **Table 1** is a summary of the number of sessions analyzed per subject, per perturbation type, as well as the average number of trials completed per session. We used all possible baseline and perturbation block trials to compute behavioral metrics and a subset of these trials for fitting factor tuning models and for fitting factor analysis models. **Table 2** displays the average number of BCI neurons that also had firing rates greater than 1Hz (assessed offline) to have sufficient samples to estimate variance. Additionally, it presents the total number of neurons used for figures in which we analyzed tuning change in individual neurons.

## Acknowledgements

This work was funded through Whitehall Grant 2022-12-071. H.M.S. was funded through the Department of Defense National Defense Science and Engineering Graduate Fellowship program. The authors would like to thank C.J. and L.W. for surgical assistance and the Biomedical Imaging Center at the University of Texas at Austin for assistance with their MRI scanner.

